# Lipid-Coated Water-in-Oil Droplets as a Passivation-Free Platform for Cost-Effective Fluorescence Spectroscopy

**DOI:** 10.64898/2026.06.26.734730

**Authors:** Joshua W Trowbridge, Aleksa Lakic, Annika Brodbeck, Dezerae Cox, Alexander F. Mason, Luke McAlary

## Abstract

Fluorescence correlation spectroscopy (FCS) provides valuable information about molecular dynamics, however, experimental setup typically requires labour-intensive passivation to prevent non-specific binding of molecules to sample containers. Furthermore, precious samples can be wasted by having to use relatively high sample volumes in existing sample containers. We overcome these major issues using a simple method of sample encapsulation into ‘water-in-oil’ droplets, using purified proteins and cell lysates as proof-of-concept. FCS of fluorescently labelled protein samples in the nanomolar (nM) range confirmed that water-in-oil droplets yield more accurate measurements than conventional open-chamber methods. We first optimized the droplet composition to prevent protein coating at the water-oil interface using pegylated-lipids. We then utilized FCS to accurately measure protein concentrations and diffusion speeds in nanolitre volumes. Additionally, we used fluorescence cross-correlation spectroscopy (FCCS) to measure enzymatic cleavage of substrate inside our droplet system, demonstrating the capacity of this platform to measure biological processes at the nanoscale. Overall, conducting FCS in droplets offers a cost-effective, robust, and accessible alternative for measuring molecular dynamics, with promising potential for high-throughput and resource-limited applications.

**TOC Image + Text:** 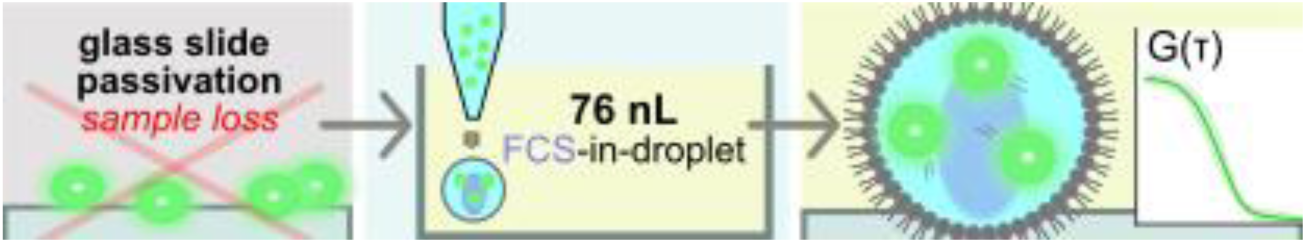

Conventional single-molecule fluorescence requires slow, expensive glass passivation procedures to prevent proteins adsorbing to surfaces. By encapsulating proteins in lipid-coated nanolitre water droplets, the passivation requirement is removed, enabling accurate measurement of protein dynamics in low nanolitre volumes. Water-in-oil droplets thus provide a passivation-free platform for fluorescence correlation spectroscopy.

## 1. Introduction

Biological systems are governed by interactions that occur across scales, with particular importance placed at the molecular scale. The study of biomolecular interactions provides useful information about biological processes for both fundamental and medical science. In particular, single-molecule fluorescence methods have been powerful for determining mechanistic details of biological processes.^[1]^ Even so, capturing protein dynamics in vitro remains challenging, requiring specialised fluorescence approaches and compatible microscopy configurations. While these methods can be performed in cells, biological noise complicates both setup and analysis. Correspondingly, most single-molecule methods utilize purified systems so that precise control over sample parameters can be achieved.^[2]^ While advantageous for discrete analysis, some major caveats arise such as loss of biological relevance and excessive sample preparation. In particular, glass slide passivation or blocking is a major requirement for most single-molecule techniques including fluorescence correlation spectroscopy (FCS) and fluorescence cross-correlation spectroscopy (FCCS).^[3,4]^

Passivation prevents the adsorption of analyte molecules to container surfaces, which would otherwise compromise data acquisition. The gold-standard passivation method uses PEG-silane functionalisation of piranha-treated glass coverslips.^[5,6]^ Not only are the chemicals dangerous, but the process can take 1-2 days and consumes expensive reagents such as biotin-PEG.^[5,6]^ Additionally, it can be a sensitive process as contamination or user-error at any stage necessitates repeating the entire protocol.^[5]^ This highlights that alternative, safer, and more cost-effective approaches are required to bypass glass passivation. Further, conventional single-molecule experiments can require relatively high sample volumes, which presents an additional obstacle when working with scarce or precious protein samples.

To address these limitations, we demonstrate the use of water-in-oil droplets as tuneable, cell-sized, nanolitre volume compartments to facilitate FCS measurements. Water-in-oil droplets have been used in many applications across biology, including to support gene expression studies^[7]^, droplet array generation^[8,9]^ and smFRET^[10,11]^. FCS has been applied to water-in-oil droplets through the characterisation of microemulsion droplets^[12]^ and quantification of droplet content and flow in microfluidics-generated droplets.^[13]^ To our knowledge, however, they have never been used as inert, ultralow-volume compartments to conduct measurements of protein dynamics. By using water-in-oil droplets to perform FCS measurements, the need for extensive glass passivation is eliminated. Instead, molecules of interest can be encapsulated within the aqueous phase of a water-in-oil droplet, and fluorescence measurements taken from within the droplets. To demonstrate the versatility of this approach, we perform FCS measurements on both purified protein and crude cell lysates. Furthermore, by adopting an approach typically used to create synthetic cell membranes,^[14,15]^ we incorporate lipids into the oil phase to minimise non-specific protein binding at the water-oil interface. Our method to generate droplets takes only minutes, rather than days of preparation. Together, these results show that lipid monolayer-coated water-in-oil droplets are a powerful and versatile platform for fluorescence analysis via FCS, and potentially other microscopy techniques, without the need for passivation.

## 2. Results and Discussion

### 2.1. PEGylated lipid water-in-oil droplets prevent non-specific protein binding

Water-in-oil droplets can provide discrete aqueous compartments in which proteins can be encapsulated, with the oil phase acting as an immiscible barrier. Considering that making such a system is as simple as mixing water with oil, we set out to understand how biomolecules would behave in these systems. Here, we used purified forms of two constitutively fluorescent proteins, mCherry (27 kDa mass, 610 nm emission) and GST-tagged Dendra2 (glutathione S-transferase-tagged 54 kDa mass, 507 nm emission when green), to assess protein behaviour. These proteins were chosen as they have different fluorescence spectra and masses, allowing for easy comparison in imaging and FCS experiments.

In simple water-in-oil droplets (aqueous phase: GST-Dendra2/mCherry in buffer, oil phase: 1:1 v/v AR20 silicone oil: hexadecane), a significant accumulation of GST-Dendra2 was observed at the water-oil interface, indicating substantial protein coating **(Figure 1A and B)**. In contrast, mCherry did not coat the water-oil interface, suggesting this phenomenon was due to a physicochemical feature of GST-Dendra2 **(Figure 1C, D and S1)**. To mitigate the protein adsorption of GST-Dendra2 at the water-oil interface, the ‘lipid-out’ technique was used,^[13]^ where lipids dissolved in the oil phase can assemble a monolayer around the aqueous droplet upon deposition, a common method used to form synthetic cell compartments.^[14,15]^ Two lipid-out oil formulations were tested to compare GST-Dendra2 coating at the water-oil interface: (i) 1,2-diphytanoyl-sn-glycero-3-phosphocholine (DPhPC) and (ii) DPhPC with 1 mol% 1,2-distearoyl-sn-glycero-3-phosphoethanolamine-N-[methoxy(polyethylene glycol)] (DSPE-mPEG) (PEGylated lipid), in the same oil mixture as before. DPhPC was chosen as it is a saturated lipid that forms stable, robust monolayers at oil-water interfaces.^[16,17]^ DSPE-mPEG was chosen as the PEG group can create a steric barrier that inhibits non-specific protein adsorption at interfaces.^[6,18]^ Line intensity profiles indicate that the inclusion of DPhPC in the oil-phase causes a reduction in GST-Dendra2 coating at the water-oil interface **(Figure 1A and B)**, suggesting that the lipid monolayer effectively reduces non-specific protein interactions. The addition of a PEGylated lipid did not produce a statistically significant difference in protein coating intensity compared to DPhPC alone **(Figure 1C,D)**, but was retained as an additional measure, given the well-characterised use of PEG-functionalisation to mitigate non-specific protein adsorption.^[19]^ Having GST-Dendra2 and mCherry in the same droplet did not change the coating properties of either protein **(Figure 1D)**. A key feature of using lipid-coated droplets is that it is not restricted to a specific lipid mixture in the oil-phase, rather this composition can be easily tuned.^[14]^ We exploited this here to fine tune the resistance of the droplet boundary to adsorption of the proteins encapsulated in the droplet. As expected, mCherry showed no coating in any oil condition **(Figure 1C)**. Although not all proteins adsorb at the water-oil interface, the lipid composition used here (DPhPC + 1 mol % mPEG-DSPE) was sufficient to enable FCS measurements of two spectrally distinct purified proteins, suggesting broad compatibility with other protein samples.

**Figure 1.**
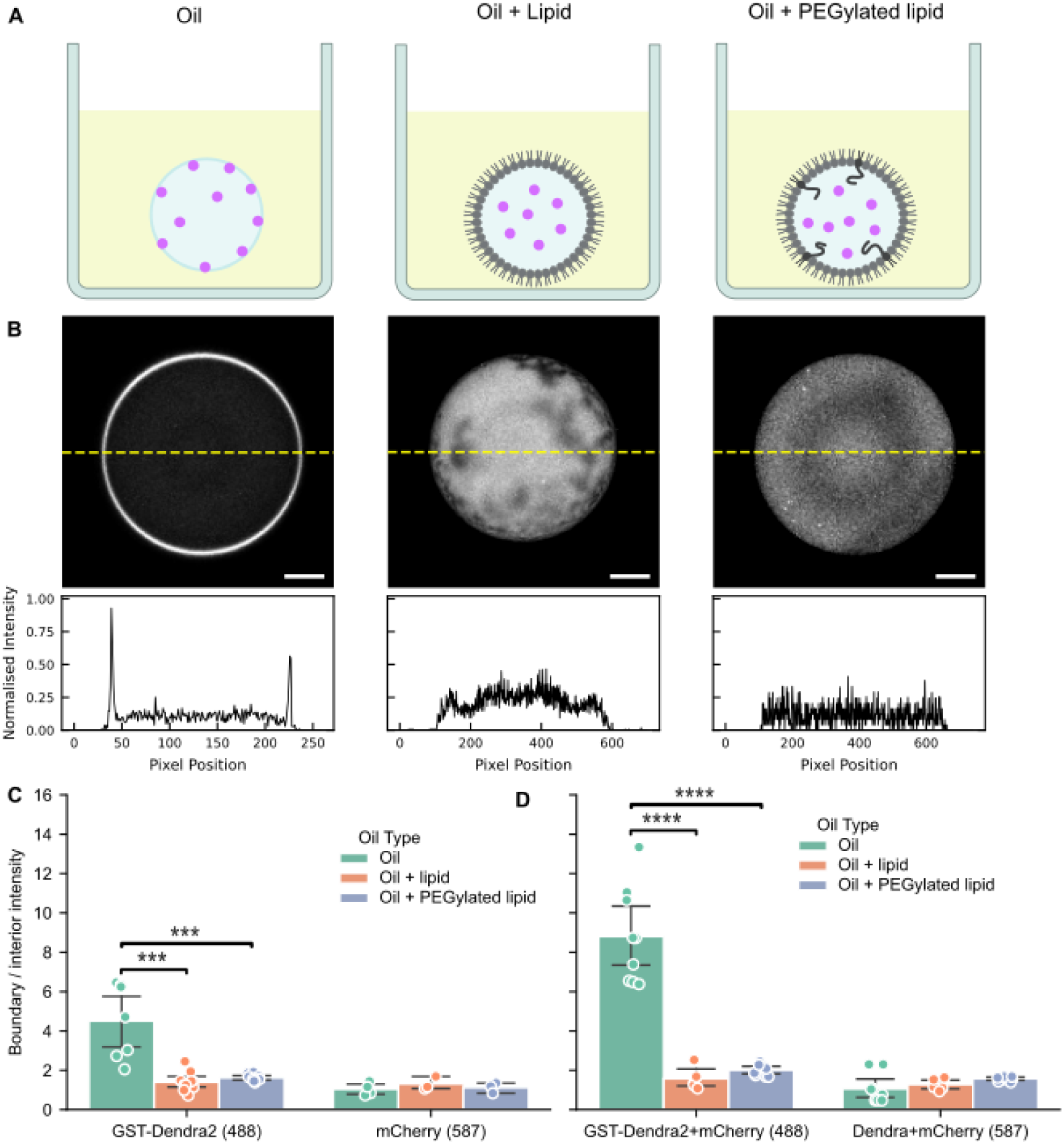
Protein coating at the water–oil interface of droplets with varying oil compositions. (A) Schematic of water-in-oil droplets for three oil compositions: oil only, oil + DPhPC (oil + lipid), and oil + DPhPC + DSPE-mPEG (oil + PEGylated lipid). (B) Representative green fluorescence images (488 nm excitation, Top) showing GST-Dendra2 coating at the water-oil interface for each condition, with corresponding line intensity profiles (Bottom) measured along the yellow dashed lines. (C, D) Quantification of protein coating at the oil water interface for droplets containing GST-Dendra2 or mCherry only, measured with 488 nm or 594 nm excitation respectively (C) or containing GST-Dendra2 and mCherry measured at both 488 nm and 594 nm excitation (D). Statistical significance was determined by one-way ANOVA within each protein group and wavelength, followed by post-hoc pairwise t-tests with Holm-Bonferroni correction for multiple comparisons. ns: p > 0.05; *: p ≤ 0.05; **: p ≤ 0.01; ***: p < 0.001, ****: p < 0.0001. n = 4–10 independent images per condition. Scale bars: 50 µm.

### 2.2 Robotic dispensing and manual pipetting both create viable water-in-oil droplets

Standard pipettes can dispense in the low microlitre range, whereas robotic systems can achieve volumes in the low nanolitre range. We have previously developed a robotic liquid-handling system called DIB-BOT, composed of a nanoinjector mounted on a 3D printer, which enables dispensing of volumes as low as 4.2 nL with improved morphological consistency and positional control.^[20]^ Built from standard 3D-printer components at a low cost, DIB-BOT provides an accessible alternative to commercial automated liquid-dispensing platforms. Here, we assess both hand-pipetting and DIB-BOT generated droplets for usage in FCS experiments of fluorescent proteins.

We generated droplets by manual pipetting (1 µL per droplet) and by the DIB-BOT (76 nL per droplet) and sought to make morphological comparisons of size and circularity (eccentricity; **Figure 2A**). Hand-pipetted droplets showed a greater variability in area and produced droplets with a higher degree of eccentricity, reflecting the inconsistencies introduced by manual handling. In contrast, DIB-BOT generated droplets were uniform in area (CV = 35.4 %, n = 21) and more circular on average than manually pipetted droplets with eccentricity values closer to zero **(Figure 2B)**. Additionally, droplets generated by manual pipetting were more prone to deformation and morphological defects. While the differences in morphology are unlikely to influence internal protein interactions, they highlight the superior reproducibility of the DIB-BOT method in maintaining consistency across assays, further supporting the advantages of automated droplet deposition for the formation of consistent synthetic compartments to facilitate FCS. Beyond reproducibility, the DIB-BOT enables dispensing of sub-100 nL, and in our case 76 nL, increasing the number of independent experiments per unit reagent by more than an order of magnitude relative to manual pipetting **(Figure 2C)**. The requirement for the DIB-BOT is not inherent to our approach, since droplets can also be deposited via pipetting at the cost of automation, reduced reaction volumes and droplet uniformity reported above.

**Figure 2:**
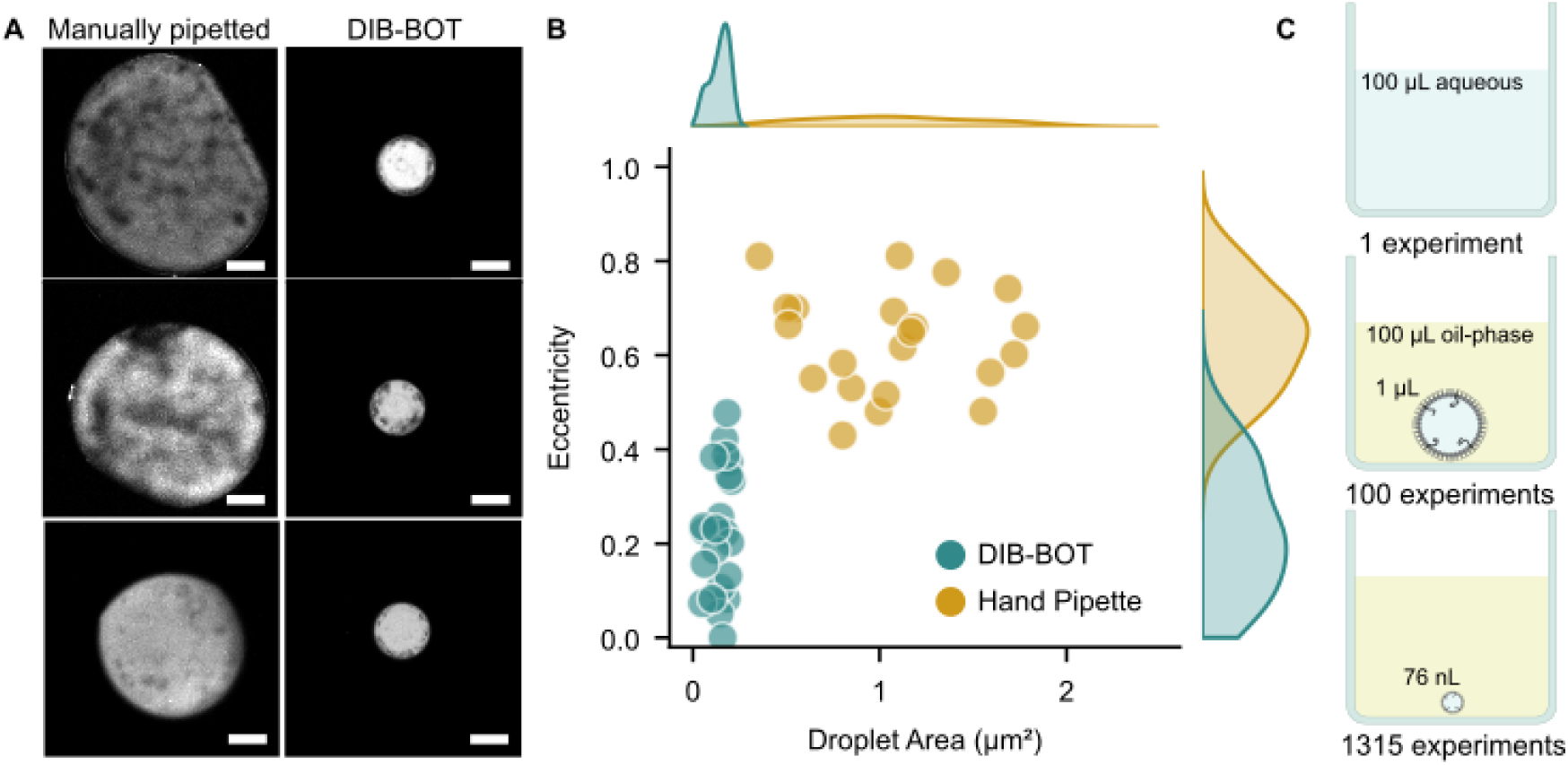
DIB-BOT dispensing compared to manual pipetting. (A) Fluorescence images of droplets produced by each method: 1 µL droplets by manual pipette (Left) and automatically dispensed via the DIB-BOT (Right). Droplets were plotted on the same scale for comparison. Scale bars: 200 µm. (B) Joint plot showing the distribution of droplet area and eccentricity for both methods (n = 21 per method). Statistical significance was determined by Kolmogorov–Smirnov test comparing droplet eccentricities only (D = 0.9545, p-value = 4.2 × 10⁻¹¹) indicating a significant difference. (C) Schematic comparison of reaction throughput per 100 µL of reagent: one bulk aqueous reaction (100 µL), ∼100 manually pipetted droplets (1 µL each), or ∼1315 DIB-BOT droplets (76 nL each).

### 2.3 Fluorescence correlation spectroscopy of protein diffusion inside droplets is accurate

Having established the droplet platform, we next assessed whether FCS could be performed accurately within lipid-coated droplets using purified proteins. We compared FCS in our droplets to standard glass-bottomed Ibidi µ-well chamber slides, with key measures being protein diffusion and concentration **(Figure 3A)**. We used purified recombinant proteins GST-Dendra2 and mCherry, both at a final concentration of 200 nM. Importantly, the glass slides did not go through time-consuming and expensive “blocking” procedures, allowing for a direct comparison between the isolated and dynamic environments in the droplets as opposed to potential interactions between proteins and the glass surface. Sulforhodamine-B (SRB) was used as a standard to calibrate FCS measurements (diffusion coefficient of 420 µm^2^/s).^[21]^ For chamber samples, we manually pipetted 100 µL of sample, whereas droplet samples were composed of 76 nL volumes produced by DIB-BOT, which is a 1300-fold decrease in sample amount. All samples gave appreciable autocorrelation curves **(Figure 3 B-C)**, indicating that both chamber and droplet samples were reporting fluorescence dynamics expected for FCS.

**Figure 3.**
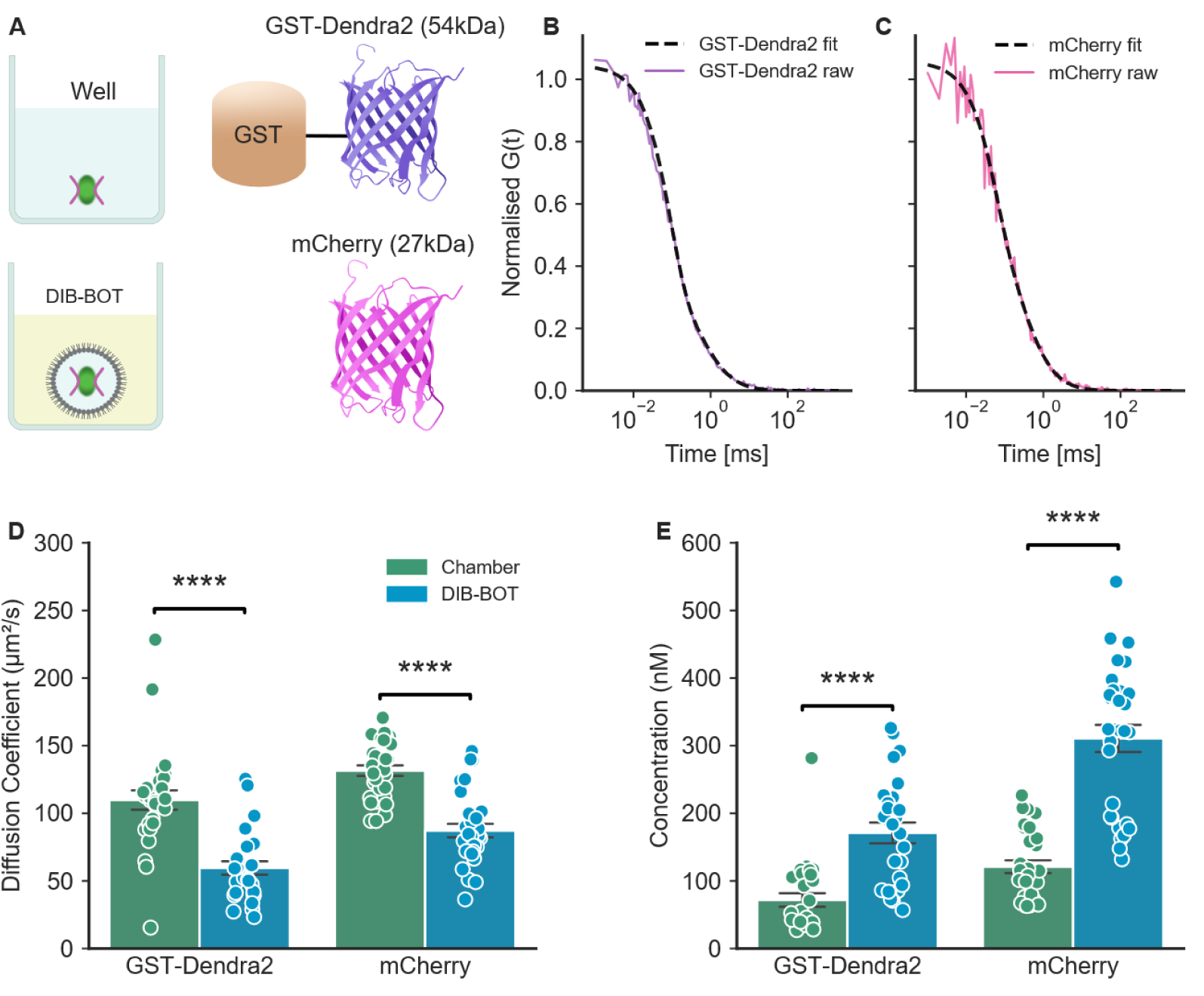
Fluorescence correlation spectroscopy (FCS) of fluorescent proteins inside droplets. (A) Schematic of FCS either occurring within 100 µL of sample in an untreated 8 chamber ibidi glass chamber slide or a 76 nL droplet within 100 µL of PEGylated lipid in an 8 well ibidi glass chamber slide. Schematic of GST-Dendra2 with the bulky GST tag and mCherry. (B-C) FCS autocorrelation curves denoting the raw data and the 3D Gaussian diffusion model fit of GST-Dendra2 (purple) and mCherry (pink) both within droplet conditions. (D-E) Diffusion coefficient (D) and concentration (E) of GST-Dendra2 and mCherry either in DIB-BOT (blue) or in chamber (green) conditions. Error bars denote standard error of the mean (SEM); n=27-30 replicates from 3 independent replicates. Statistical comparison between chamber and DIB-BOT conditions were made using Student’s or Welch’s t test following Levene’s test for equality of variances p < 0.0001; ****.

Autocorrelation curves were fitted using a 3D Gaussian diffusion triplet model ^[22–24]^. The collated autocorrelation fits were consistent over independent replicates **(Figure S2)**. These fits from autocorrelation curves returned discrete values for diffusion coefficients **(Figure 3 D)** and concentrations **(Figure 3 E)**. For mCherry in droplets, we measured a diffusion coefficient of 87.20 ± 27.22 µm^2^/s (mean ± SD, n = 30), which is consistent with previous reports of the similarly sized green fluorescent protein (GFP, 27 kDa) in aqueous buffer (80 – 100 µm^2^/s).^[25,26]^ The diffusion of mCherry in the chamber was measured at 131.53 ± 21.23 µm^2^/s (mean ± SD, n = 30), which is faster than expected. Meanwhile, GST-Dendra2 in droplets had a diffusion of 59.54 ± 25.54 µm^2^/s (mean ± SD, n = 27), which is consistent with its theoretical diffusion of 67 µm^2^/s (derived from the Stokes-Einstein equation) and corresponds to its increased mass (54 kDa) compared to mCherry (27 kDa). The diffusion coefficient of GST-Dendra2 in the chamber was 109.78 ± 38.00 µm^2^/s (mean ± SD, n = 28), unexpectedly close to that of the smaller mCherry (131.52 µm²/s), despite their substantial difference in molecular mass. Importantly, we found that the measured diffusion speeds for our model proteins when measured in droplets were closer to theoretical measurements when compared to chamber samples. While the exact cause is unclear, a likely reason is due to an increase in temperature of the microscope body or objective lens, a known cause of elevated protein diffusion rates^[27–29]^. This effect was absent in the droplet measurements, suggesting that the oil reservoir may provide sufficient thermal mass to buffer the droplet against temperature changes, resulting in more accurate diffusion readings, however this hypothesis remains to be tested directly.

Meanwhile, measurements of protein concentrations were consistent within technical replicates for both chamber samples and droplets, but there was some deviation between experimental replicates. Overall, measured concentrations of GST-Dendra2 were closer to the expected 200 nM range for droplet (170.96 ± 80.19 nM (Mean ± SD n = 27)) than compared to chamber samples (71.80 ± 53.23 nM (Mean ± SD n = 28)). However, for mCherry, the chamber sample (121.02 ± 51.60 nM (Mean ± SD n = 30)) was more consistent with the 200 nM concentration when compared to the droplet sample (310.74 ± 109.87 nM (Mean ± SD n = 30)). Both concentrations measured in the chamber were lower than their droplet counterpart, indicating there is protein adsorption at the glass interface. Deviations from expected concentrations of 200 nM occur likely due to adsorption when diluting concentrated samples and from differences in how UV/Vis spectroscopy and FCS both determine concentration. Despite this, the values are close to what was expected, showing that our droplet method is highly appropriate for FCS to determine analyte concentration and diffusion.

### 2.4 Protease cleavage can be measured inside droplets using fluorescence cross-correlation spectroscopy

Having confirmed the robustness of our droplets for purified samples with FCS, we next challenged the system with a more technically demanding measurement in a more complex biological sample. To this end, we elected to perform fluorescence cross-correlation spectroscopy (FCCS) in crude mammalian cell lysates. In brief, FCCS can be used to measure the interaction between two fluorescently labelled molecules that emit at different wavelengths by correlating instances where both fluorescent molecules inhabit the detection volume simultaneously. While FCCS requires rigorous controls, it enables extremely sensitive, time-resolved measurement of biologically relevant molecular processes in homogenous mixtures such as crude lysates.

We cultured mammalian 293T cells and transfected them with synthetic constructs that fuse acGFP1 to mCherry with a linker sensitive to tobacco etch virus protease (called acGFP1-TEV-mCherry) or a normal linker (acGFP1-mCherry), allowing us to monitor protease cleavage with FCCS. Controls were cells expressing either acGFP1 or mCherry alone. Cells were lysed via gentle detergent, clarified via centrifugation and the soluble component used for droplet generation. Sulforhodamine B was again used as an internal standard to calibrate autocorrelation curves of all samples, and the final volumes used were 76 nL for each sample. Importantly, co-encapsulated but unfused acGFP1 and mCherry in crude lysate showed no cross-correlation, confirming that both species must be covalently bound to produce a cross-correlation signal **(Figure 4B)**. In contrast, we observed strong cross-correlation in samples containing acGFP1-mCherry or acGFP1-TEV-mCherry, indicating both GFP and mCherry were diffusing through the measurement volume simultaneously.

**Figure 4.**
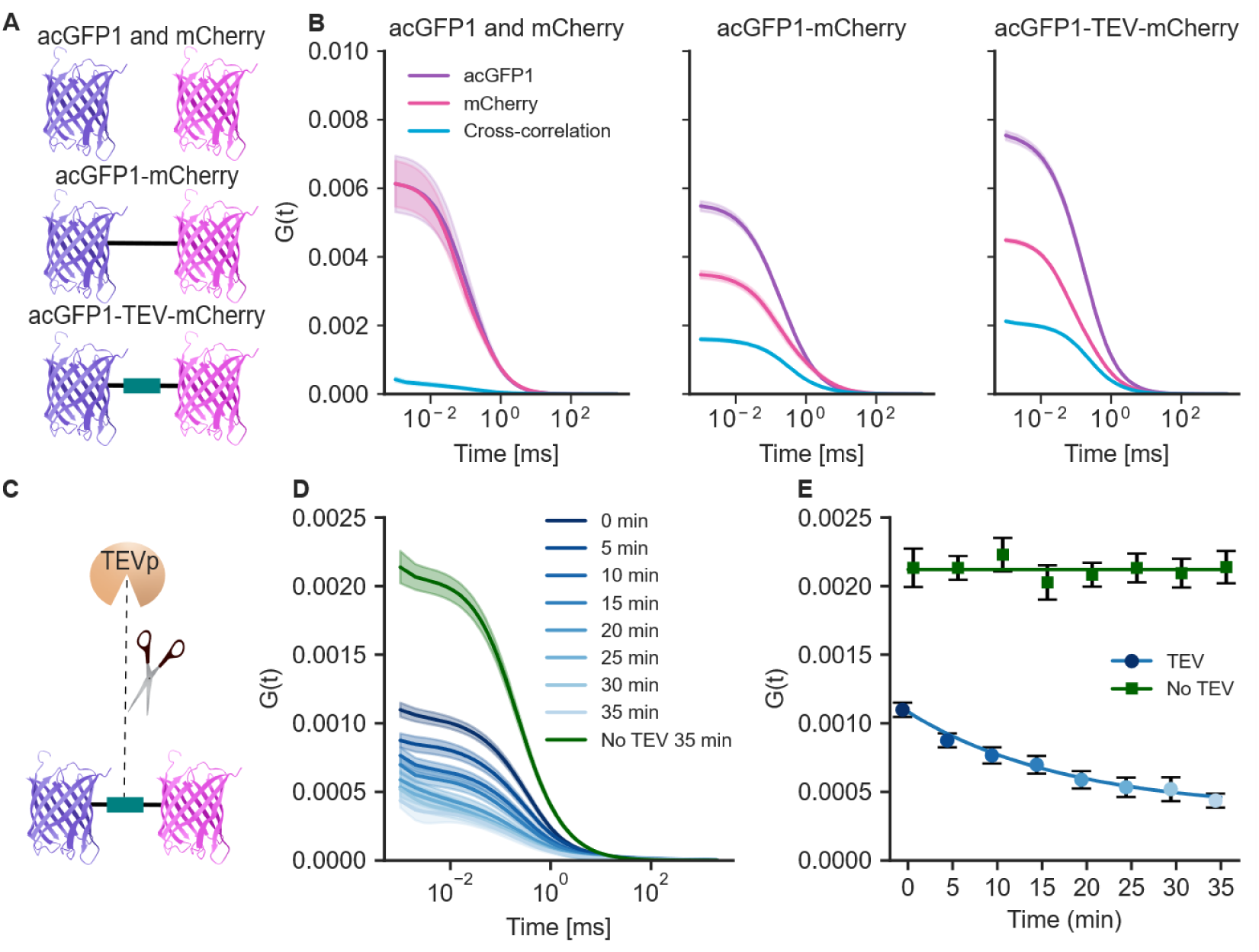
Using fluorescence cross-correlation spectroscopy (FCCS) to measure protease cleavage in droplets. (A) Schematic depicts the three constructs used, acGFP1 and mCherry individually, acGFP1-mCherry, and acGFP1-TEV-mCherry. (B) FCCS autocorrelation curves of acGFP1 (490 nm; purple) and mCherry (587 nm; pink) across three different constructs. Blue curve denotes instances of cross correlation. Error bars represent SEM; n=15-16 from 2 independent replicates. (C) Depiction of TEV protease cleaving acGFP1-TEV-mCherry. (D) Cross-correlation curves of acGFP1-TEV-mCh with TEV protease present at 1:75 ratio over 35 minutes from dark blue at 0 minutes to light blue at 35 minutes. Cross-correlation curve of acGFP1-TEV-mCherry no TEV protease in green. (E) Initial cross-correlation values over time for acGFP1-TEV-mCherry with (blue) or without (green) TEV protease. An exponential decay function was fitted to the with TEV protease sample over 35 minutes to quantify cleavage kinetics and a zeroth-order polynomial fit has been added to without TEV protease sample to account for baseline cross-correlation. Error bars represent SEM; n=26-30 per individual timepoint from 3 independent replicates.

We next used FCCS to monitor an enzymatic reaction in real time within the droplet platform. The acGFP1-TEV-mCherry construct enables direct measurement of TEV protease cleavage activity using FCCS which has been described in the literature ^[30]^. Theoretically, this separation would lead to a measurable decrease in cross-correlation across time. We first confirmed that our purified TEV protease was active and that the acGFP1-TEV-mCherry could be cleaved via immunoblot **(Figure S3)**. We then established a time-course assay in crude cell lysate, where the fluorescent fusion constructs (acGFP1-TEV-mCherry or acGFP1-mCherry as a negative control) were mixed with TEV protease prior to droplet creation via the DIB-BOT. Upon generation of the droplet, the sample was quickly transferred to the microscope with minimal dead time (∼5 min). Across the time course, we observed a decrease in the cross-correlation in acGFP1-TEV-mCherry droplets containing TEV protease, but not when TEV protease was absent from the reaction **(Figure 4D and E)**. As expected, the TEV protease had no effect on the cross-correlation of the acGFP1-mCherry construct **(Figure S4)**. Due to our reaction volume being only 76 nL, we only required a small volume of reactants to perform this assay, a key benefit of this approach, especially with scarce or precious protein samples. Furthermore, the DIB-BOT droplet allowed us to measure enzymatic cleavage in a consistent manner.

Despite the promising performance of using droplets to bypass slide passivation for conducting FCS measurements in solution, we have identified some limitations. Locating a small droplet under high magnification can be time-consuming, increasing the dead time between droplet deposition and image acquisition. We recommend using small wells to decrease the search space. Further, we have not assessed compatibility with long-term imaging, as at most our acquisition time was 35 minutes for the TEV protease assay. This work validated the droplet platform specifically for FCS/FCCS, though the approach may be compatible with other techniques for studying protein dynamics. Finally, realising the true benefits of reproducibility and reduced reaction volumes requires access to an auto-dispensing machine like the DIB-BOT, however we do note that hand pipetting is still accessible for creating contained ∼ 1µL sample volumes. Furthermore, our water-in-oil droplet approach addresses a broader need for accessible single-molecule microscopy that circumvents costly and time-consuming glass passivation procedures while conserving limited sample volume **(Tables S1-S3)**. We estimate that the method developed here costs roughly $3.30 AUD per sample in reagents **(Table S3)** and that the preparation time prior to sample measurement is reduced to 17 h (shorter if not doing overnight lipid film drying) compared to a standard passivation method **(Table S1)** [5] and comparable to an intensive but rapid method **(Table S2)** [6]. Notably, our method does not require the use of the highly dangerous piranha solution, thus greatly increasing user safety.

Beyond FCS/FCCS, the lipid-coated droplet platform should be compatible with other solution-based fluorescence techniques that are affected by non-specific binding to surfaces, including smFRET or single-particle tracking, and with potential applications testing protein interactions on disease related targets, enzymatic assays, and drug screening against protein targets, where reduced sample volumes and passivation-free methods are beneficial. Combined with the DIB-BOT, the platform further reduces sample requirements to ultralow volumes compared to the lower limit of a regular pipette, increases throughput, and there is also a potential for automating content mixing, which was not explored here.

## 3. Conclusion

We have demonstrated that PEGylated-lipid-coated water-in-oil droplets provide a rapid, cost-effective, resource-conscious and accessible alternative to conventional glass passivation for single-molecule fluorescence measurements of proteins. FCS and FCCS measurements of purified proteins and cell lysates in droplets yielded accurate diffusion coefficients and enabled real-time monitoring of enzymatic activity, all within sub-100 nanolitre volumes. Notably, the droplet-based approach reduces the sample volume required for a single FCS experiment by more than two orders of magnitude compared to conventional methods. Combined with automated droplet generation via DIB-BOT and the tunability of the lipid interface, this platform is well-positioned as a versatile tool for measuring biomolecular biophysics.

## 4. Experimental Section

*Experimental Subheading*: Experimental Details.

### 4.1 Materials

Glass capillaries (3-000-203-G, 1.14 mm O.D. x 3.5″ length, Drummond) and NanojectII (Drummond) were purchased from Adelab Scientific. DPhPC, DSPE-mPEG, hexadecane, AR-20 silicone oil and chloroform were purchased from Sigma-Aldrich.

### 4.2 Oil/lipid prep

Lipid stocks in chloroform were aliquoted into glass vials and dried down to a thin film under a gentle stream of nitrogen gas, prior to storing under vacuum for a minimum of 3 h (typically overnight) at room temperature in a vacuum desiccator to remove residual solvent. Dried films were stored under nitrogen at − 20 °C if not used immediately.

Lipid films were resuspended in a premixed oil solution consisting of silicone oil (AR-20) and hexadecane in a ratio of 1:1 v/v to a concentration of 5 mg mL^−1^. The lipid-in-oil solution was vortexed briefly and then bath-sonicated for 30 min at 25–35 °C before use.

Three oil formulations were prepared in AR-20 silicone oil/hexadecane (1:1 v/v): oil without lipid, 5 mg mL⁻¹ DPhPC (oil + lipid), and 5 mg mL⁻¹ DPhPC:DSPE-mPEG at 99:1 mol/mol (oil + PEGylated lipid).

### 4.3 Method for water-in-oil protein droplet formation

Wells of an 8-well chambered coverglass were each filled with 100 µL of one of the three oil solutions. Inner aqueous solutions were typically prepared in 20 µL volumes in plastic PCR tubes and contained fluorescent proteins (GST-Dendra2, mCherry, or others as specified) in PBS buffer (50 mM, pH 7.4) at the concentrations stated in Section 4.7.

Aqueous solutions were deposited into the oil phase either by hand-pipetting, or by micro-injection using a NanojectII. For hand-pipetted droplets, 1 µL aqueous volumes were deposited directly into the oil using a P1 pipette. For the microinjected droplets, semi-automated deposition was performed using the DIB-BOT as described previously.^[20]^ Briefly, a pre-pulled microcapillary (Drummond, 3-000-203-G) was mounted to the NanojectII, backfilled with hexadecane, and filled with aqueous solution. Droplets were injected directly into the oil-filled chamber at nL volumes, with injection volume set using the binary switches on the NanojectII control unit to deliver 76 nL per droplet.

### 4.4 Bacterial expression and purification of proteins

Bacterial expression plasmids used in this study included pRK793 from David Waugh (Addgene: #8827, expression of TEV protease^[31]^), pET-MBP-mCherry from Scott Gradia (Addgene: #29747, expression of mCherry), and pGEX6P-1-Dendra2 from Periklis Pantazis (Addgene: #82436, expression of GST-Dendra2^[32]^). All plasmids were received as stabs and miniprepped to produce pure DNA for transformation into BL21(DE3) cells via heatshock.

TEV protease was expressed and purified according to previous work.^[31]^ Autoinduction was used to express both mCherry and GST-Dendra2.^[33]^ Briefly, overnight starter cultures were made in LB and shaken at 200 rpm. Main cultures of terrific broth (TB) supplemented with sugars (glucose 40 mg/L, glycerol 500 mg/L, α-lactose monohydrate 200 mg/L), salts (NH_4_Cl 526 mg/L, KH_2_PO_4_ 680 mg/L, Na_2_HPO_4_ 710 mg/mL, Na_2_SO_4_ 142 mg/L), and antibiotic (GST-Dendra2 = 100 µg/mL carbenicillin, mCherry = 50 µg/mL kanamycin), were inoculated with starter cultures and shaken at 220 rpm for 6 h at 37 °C, after which they were shaken at 20 °C for 24 h. Expression cultures were harvested via centrifugation at 6000 ×*g* for 5 min, after which the pellet was frozen overnight at -20 °C.

Pellets of GST-Dendra2 were resuspended into ice-cold lysis buffer (1× PBS supplemented with 5 mM EDTA and 1 mM PMSF, pH 7.4) and sonicated using a disruptor horn (40% amplitude, 20 sec on, 20 sec off ×5) to lyse bacteria. Lysates were then clarified via centrifugation at 40000 × *g* for 20 min and the clarified lysate was added to glutathione-resin and incubated at 4 °C with rocking overnight to promote binding. Resin was washed with 5 volumes of ice-cold lysis buffer 5 times and then GST-Dendra2 protein was eluted in 1× PBS with 5 mM glutathione in 2 elutions. Protein purity was determined via SDS-PAGE and concentration determined using the Dendra2 extinction coefficient at 490 nm excitation (45000 M^-1^ cm^-1^).^[34]^ Samples of GST-Dendra2 were snap frozen and stored at -80 °C.

Pellets of MBP-mCherry were resuspended into ice-cold lysis buffer (20 mM Tris, 30 mM imidazole, 500 mM NaCl, 1 mM PMSF, 1× EDTA-free protease tablet, pH 8.0) and sonicated as above to lyse bacteria. Lysates were clarified via centrifugation at 40000 × *g* for 20 min, filtered through a 0.22 µm filter and applied to a HisTrap column equilibrated in binding buffer (20 mM Tris, 30 mM imidazole, 500 mM NaCl, pH 8.0). Following binding, the HisTrap column was washed with 200 mL of binding buffer and elution was performed over 50 mL with a gradient of 0-100% elution buffer (20 mM Tris, 500 mM imidazole, 500 mM NaCl, pH 8.0). Fractions containing pure MBP-mCherry were pooled and incubated overnight in binding buffer with the addition of 1 mg/mL TEV protease to cleave the MBP tag from the mCherry protein. Protein samples were then concentrated via centrifugal filter units, filtered through a 0.22 µm filter and applied to a S75 size-exclusion column equilibrated in 1× PBS to separate MBP (43 kDa) from mCherry (27 kDa). Fractions containing pure mCherry were pooled and the concentration determined using the mCherry extinction coefficient at 587 nm (72000 M^-1^ cm^-1^) prior to snap freezing and storage at -80 °C.

### 4.5 Mammalian tissue culture, plasmids, transfections, lysis, and immunoblot

Plasmids for mammalian tissue culture included AcGFP-N1 from Michael Davidson (Addgene: #54705, expression of AcGFP), pmCherry-C1 from Takara Bio, pcDNA3.1(+) AcGFP-mCherry (designed in-house and generated by Genscript), and pcDNA3.1(+) AcGFP-TEV-mCherry (designed in-house and generated by Genscript). All constructs were transformed into DH5α cells and miniprepped to produce pure plasmid.

Mammalian HEK293T cells were purchased from CellBank Australia and verified via STR profiling. HEK293T cells were cultured in T25 flasks in growth medium (DMEM high-glucose, 1× glutamax, 10% FBS) at 37 °C with 5% atmospheric CO_2_. Cells were routinely passaged at 70% confluency via trypsin-EDTA treatment. Prior to transfection, cells were plated at 30% confluency into 24-well plates and incubated 24 h. Lipofectamine3000 (Thermofisher) was used to transfect cells according to the manufacturer’s instructions with 500 ng of plasmid per well. Transfected cells were incubated for 48 h prior to lysate preparation.

For fluorescence measurements cells were lysed in ice-cold lysis buffer (1× PBS, 0.1% (v/v) Triton X-100, 1× DNase, 5 mM MgCl_2_, 1× Halt Protease inhibitor). Insoluble material was pelleted via centrifugation at 17000 ×*g* for 30 min, after which the supernatant was collected and frozen at -20 °C.

Cleavage of the AcGFP-TEV-reporter was confirmed via immunoblot. For this purpose, cells were harvested, spun at 10,000 ×g to pellet, and lysed in 200 µL ice-cold lysis buffer. Lysates were sonicated to clear DNA. To assess cleavage activity, purified TEVp was added to the lysate and incubated for 1 h at room temperature. The whole cell lysates were mixed with sodium dodecyl sulfate (SDS) sample buffer in a 1:3 ratio. Aliquots (15 µL) of lysates were separated via SDS-PAGE for approximately 1 h at 150 V in Mini-PROTEAN precast gel stain-free any kDa gels (BioRad). Proteins were transferred to a polyvinylidene difluoride (PVDF) membrane using western transfer buffer for 1 h at 100 V. Blots were blocked in 5% skim milk in tris-buffered saline and tween solution (TBST) buffer for 1 h at room temperature and probed overnight at 4 °C with rabbit anti-GFP (1:5000, ab290, Abcam). The next day blots were washed 3 × 5 min in TBST and probed for 1 h at room temperature with goat anti-rabbit HRP-conjugated secondary antibody (1:5000, Dako P044801-2). Then blots were washed again for 3 × 5 min in TBST, visualized by enhanced chemiluminescence (BioRAD ClarityTM Western ECL Substrate) and imaged on ChemiDocTM MP using chemiluminescent settings.

### 4.6 Image acquisition of droplets

A Leica SP8 falcon microscope (Wetzlar, Germany) was used for acquiring images of both hand pipette and DIB-BOT droplets. The white light laser was turned on at a power of 85% and 80 MHz, the 10× objective was used and images were taken at 16 bits with a pixel format of 1264 × 1264 and a 100 Hz scan speed.

### 4.7 Fluorescence correlation spectroscopy (FCS) and fluorescence cross-correlation spectroscopy (FCCS) acquisition

Ibidi µ-slide 8 chamber glass bottom coverslips (Gräfelfing, Munich, Germany) were not treated and loaded with Sulforhodamine-B (SRB), purified GST-Dendra2/mCherry, or 293T cell lysates containing acGFP1 and mCherry, acGFP1-mCherry or acGFP1-TEV-mCherry. Samples were left to equilibrate to 25 °C and diluted to 200 nM except for cell lysates which were loaded at a 4× dilution in PBS (50 mM, pH 7.4, 200 µL total for chamber or 5 µL total for droplet) and samples were mixed directly prior to droplet preparation and imaged immediately after. FCS measurements were performed using a Leica SP8 falcon microscope (Wetzlar, Germany) using the FCS application with an 86× water immersion objective with a motorised correction collar. The white light laser was turned on at a power of 85% and 80 MHz repetition. An excitation/emission wavelength of 488/498 nm for GST-Dendra2 and acGFP1, 561/571 nm for SRB and 594/604 nm for mCherry both purified and in lysate for subsequent experiments. FCS data were collected for 10 s per technical replicate (25× for each biological replicate, n = 2-3). FCS fits were calculated using a 3D Gaussian diffusion triplet model (Equation 1) and diffusion coefficients were used to characterise the movement of samples through the medium. SRB (200 nM) was used for dye calibration as it is a derivative of rhodamine B which belongs to the xanthene family with a known diffusion coefficient of 420 µm^2^/s in water at 25 °C.^[21,35]^ For enzymatic cleavage experiments, protein was mixed with TEV protease at a 1:75 ratio and imaged as described with measurements taken manually every 5 minutes.

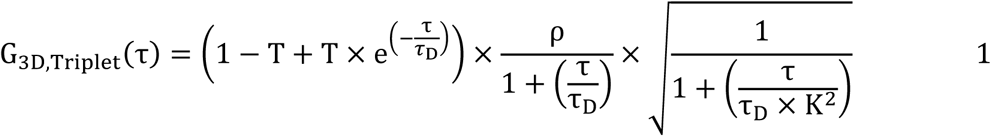

### 4.8 Analysis of droplet boundary and interior fluorescence

Semi-automated analyses of droplet images were carried out using custom Python scripts. Briefly, droplets were segmented from fluorescence images by ellipse fitting to thresholded intensity, producing labelled masks for each image. Poorly fitted masks were manually corrected. For each droplet, mean and maximum fluorescence intensities were extracted for the boundary and interior regions, and boundary-to-interior ratios were computed per droplet **(Figure S5)**. Statistical significance was determined by one-way ANOVA within each protein group and wavelength, followed by post-hoc pairwise t-tests with Holm-Bonferroni correction for multiple comparisons. AI models (Claude, GitHub Co-Pilot) were used to refine analysis scripts that were generated. The analysis process was validated via manual interrogation of input and output at every stage of the pipeline. The authors take full responsibility for the integrity of the analysis script.

### 4.9 Analysis of FCS and FCCS

Raw FCS and FCCS measurements were exported as excel files and autocorrelation fits, diffusion coefficients (µm^2^/s) and concentrations (nM) were processed using custom Python scripts. These scripts were initially generated using Claude Code (Sonnet 4.6) to transfer data between programs. On the Leica SP8 falcon microscope the 3D gaussian fit was applied to the raw curves in the software prior to being exported as an excel file. Once imported into python, for a single selected replicate, raw curves and their fit were normalised to the mean of the 10 first early plateau values and plotted on a logarithmic time axis. From the software the table values which include diffusion coefficient and concentration were also exported and given a molecule identity (GST-Dendra2 or mCherry) and condition (DIB-BOT or chamber). Any technical replicate under 5 µm²/s was excluded due to the likelihood of being an artefact. A bar and strip plot were then overlaid grouped by molecule and condition. Error bars represent standard error of the mean. A Levene’s test was conducted to test variance between conditions, and then a Students or Welch’s t-test was used as needed. Significance labels were added to any instance of statistical significance (p<0.05).

For FCCS the excel files containing raw data and fit channels were processed and saved as a melted excel file containing only fit channels for each sample (acGFP1 and mCherry, acGFP1-mCherry and acGFP1-TEV-mCherry). These cleaned excel files were used to collate fits of each sample and plotted on a y axis of G(t) and a logarithmic x-axis. These cleaned excel files were then used to plot cross correlation curves over time and as initial points. Initial points were fitted with an exponential decay function for TEV protease positive samples and a zeroth-order polynomial fit for TEV protease negative samples. Error bars represent standard error of the mean.

## Acknowledgements

We would like to acknowledge the assistance of Madison Antonia Bush for drawing schematics for some figures. The authors acknowledge the facilities, and the scientific and technical assistance of the Fluorescence Analysis Facility at Molecular Horizons, University of Wollongong. D.C. was supported by an Australian Research Council Discovery Early Career Researcher Award [DE240100707], and the National Health and Medical Research Council (ROR https://ror.org/011kf5r70) [2025780]. A.F.M was supported by an Australian Research Council Discovery Early Career Researcher Award [DE230100684]. L.M. acknowledges funding from La Caixa Health, FightMND, and the NHMRC. AI was used to iteratively assist with proof-reading text for spelling and grammar as well as to generate general feedback. All suggestions made by AI during manuscript drafting (Claude Sonnet 4.6, Git-Hub Co-Pilot GPT5-mini) were reviewed by the authors before incorporation into the manuscript, and the authors take full responsibility for the final content of the manuscript.

## CRediT Statement

**Joshua W Trowbridge:** Methodology, Software, Validation, Formal Analysis, Investigation, Data Curation, Writing - Original Draft, Visualization. **Aleksa Lakic:** Methodology, Software, Validation, Formal Analysis, Investigation, Data Curation, Writing - Original Draft, Visualization. **Annika Brodbeck:** Methodology, Formal Analysis, Investigation, Writing-Reviewing and Editing. **Dezerae Cox:** Software, Visualisation, Formal Analysis, Data Curation, Writing - Review & Editing. **Alexander Mason:** Conceptualization, Methodology, Software, Visualisation, Writing - Review & Editing, Supervision, Project administration, Funding acquisition. **Luke McAlary:** Conceptualization, Methodology, Validation, Investigation, Writing - Review & Editing, Supervision, Project administration, Funding acquisition.

## Data Availability Statement

The data that support the findings of this study are available from the corresponding authors upon reasonable request. Custom python scripts for the analysis are available from Zenodo (10.5281/zenodo.20741114, 10.5281/zenodo.20741129).

