## Supplementary Information for "Lipid-Coated Water-in-Oil Droplets as a Passivation-Free Platform for Cost-Effective Fluorescence Spectroscopy"

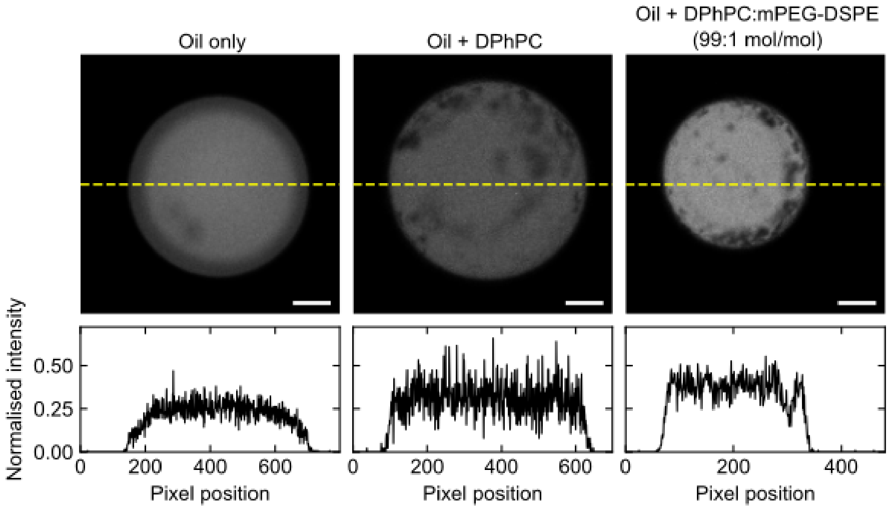


**Figure S1.** Representative yellow fluorescence images (567 nm excitation, Top) showing droplets containing mCherry for each condition, with corresponding line intensity profiles (Bottom) measured along the yellow dashed lines.

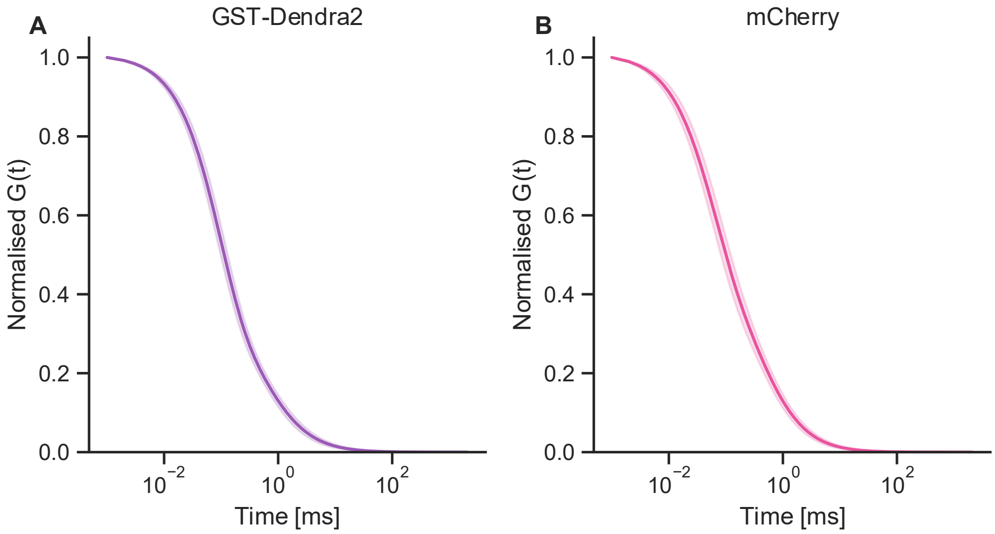


**Figure S2.** Collated 3D gaussian diffusion with triplet fits of purified protein (A) GST-Dendra2 and (B) mCherry. Error bars represent standard deviation; n = 26-30 from 3 independent replicates.

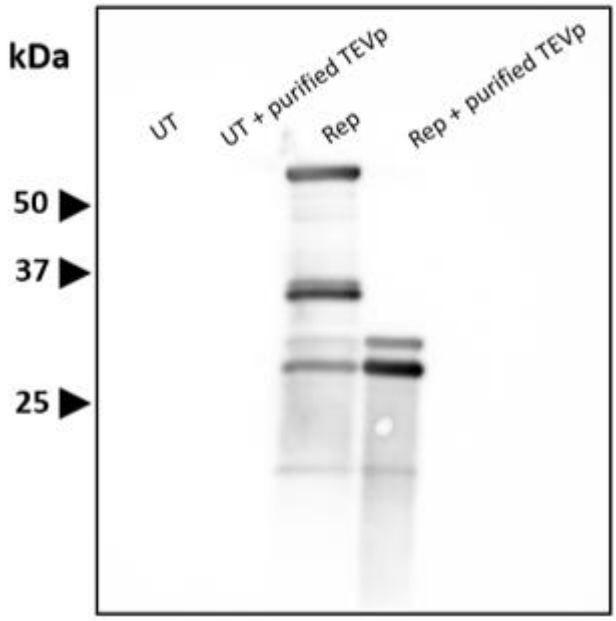


**Figure S3.** Immunoblot of AcGFP-TEV-mCherry cleavage via TEV protease. UT is an untransfected cells and Rep is the AcGFP-TEV-mCherry reporter. Some cleavage products are evident in the lysate without TEV protease likely due to proteolysis of the construct by normal cell metabolism.

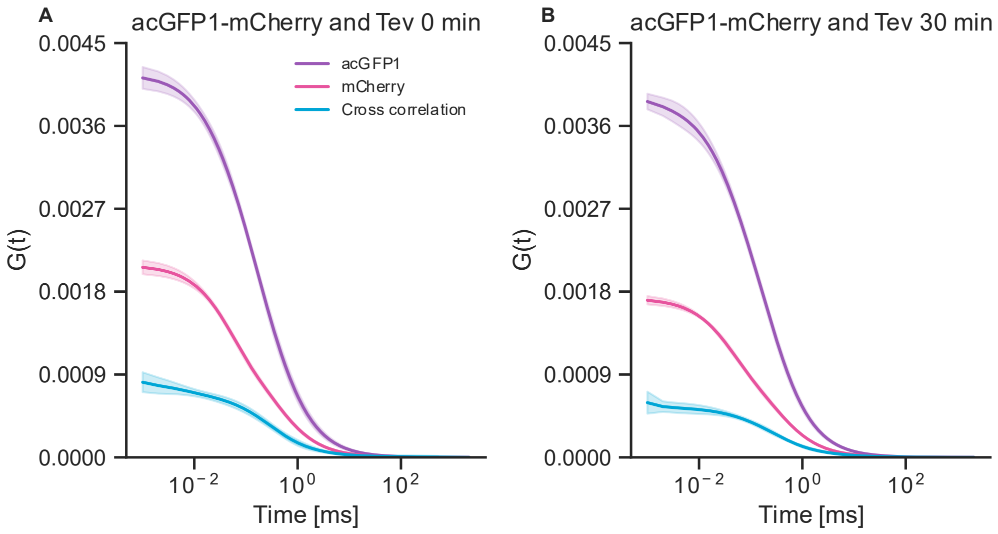


**Figure S4.** acGFP1-mCherry mixed with TEV at a 1:75 mix and read at 0 min and 30 min. Error bars represent standard deviation; n = 8 from 1 independent replicate.

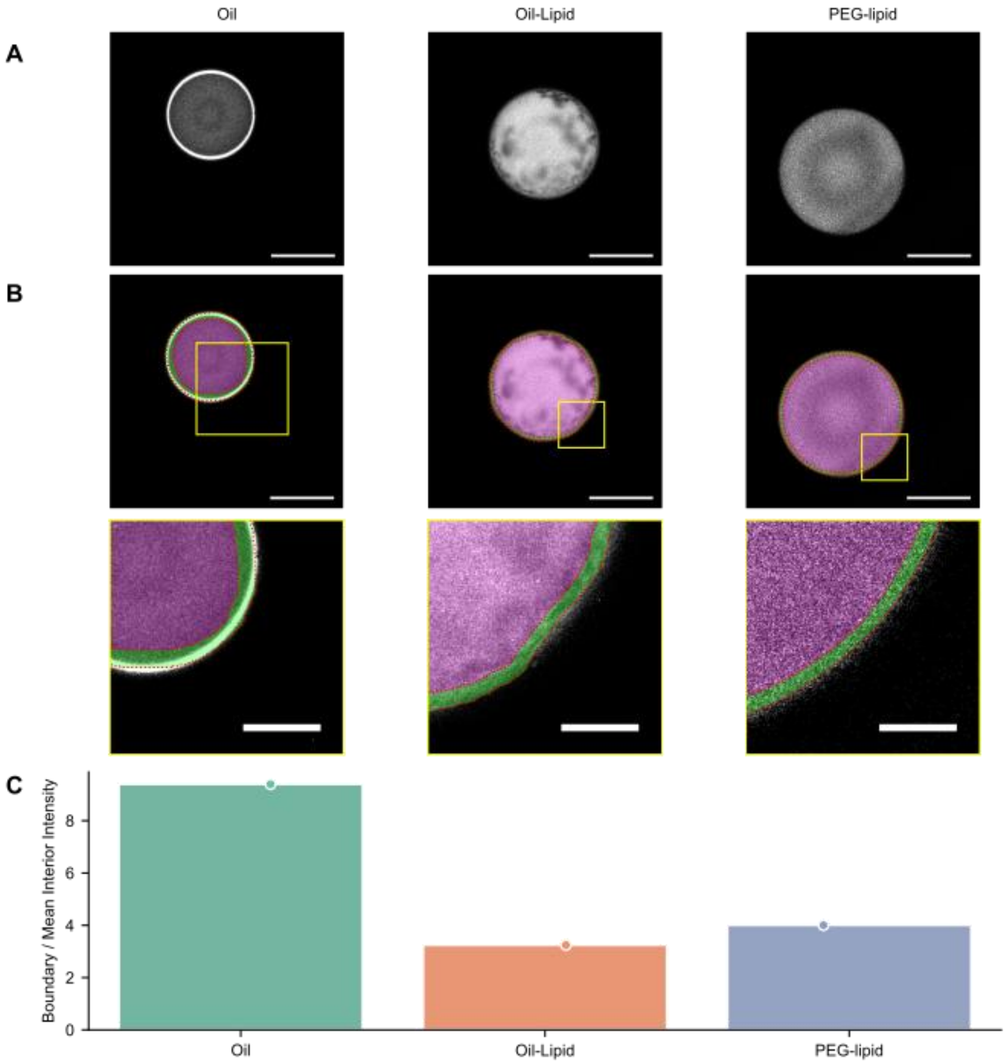


**Figure S5.** Boundary/ intensity calculation. (A) Raw, green fluorescence images (488 nm excitation) showing droplets containing GST-Dendra2 for each condition (oil, oil + *lipid, oil + PEGylated lipid* ) Scale bars: 100 µm. (B) Visualisation of thresholding approach. Boundary and interior pixels were automatically segmented from fluorescence images by ellipse fitting to thresholded intensity, producing labelled masks for each image. Boundary and interior outlines (red, dotted line), with the boundary region (green) and interior region (magenta) shaded for each respective image (Top, Scale bars: 250 µm). Zoomed regions of thresholds for each respective image (Bottom, Scale bars: 25 µm (C) Exemplar boundary/ interior intensity ratio derived from pixel intensity values of the thresholded regions in B.

**Table S1. Cost, timing and safety of standard single-molecule surface passivation.**

| Step | Reagents | Time | Cost | Risk |
|---|---|---|---|---|
| **1. Drilling Quartz Slides** | Diamond drill bit | 10 min | Per bit - $10 | Low, potential for quarts shard |
| **2. Cleaning with H2O** | MilliQ | 5 min | $0 | None |
| **3. Cleaning with Acetone** | Acetone | 20 min | 500 mL - **$72.50** | Moderate – flammable; no ignition sources |
| **4. KOH Cleaning** | KOH | 16 h | KOH 25g - **$52.80** | Low, Corrosive base |
| **5. Piranha Etching** | H2SO4 (sulfuric acid), H2O2 (hydrogen peroxide) | 20 min | Sulfuric acid 500 mL - **$136.00**<br>hydrogen peroxide 100 mL **- $125.00** | EXTREME — spontaneously heats >90C, violently oxidising, explosive with organics, fume hood required |
| **6. Coverslip Cleaning (KOH)** | 1 M KOH, MilliQ H2O | 20 min | Same as step 4 | Low, Corrosive base |
| **7. Amino-silanization** | Methanol, acetic acid, APTES (3-aminopropyl trimethoxysilane) | 30 min | Methanol 1L- **$61.90**<br>Acetic acid 500 mL- **$89.30**<br>Aptes 100 mL - **$187.00** | Moderate, APTES is toxic/neurotoxic irritant; methanol flammable |
| **8. PEGylation Round 1** | NHS-ester mPEG, biotinylated NHS-ester PEG, sodium bicarbonate buffer | 16 h | NHS-ester mPEG 1g - **$269.11**<br>biotinylated NHS-ester PEG - **$384.43**<br>sodium bicarbonate 500g - **$86.40** | Low; NHS-ester moisture-sensitive — store frozen under N2 |
| **9. PEGylation Round 2** | MS(PEG)4-short NHS-ester PEG (333 Da), sodium bicarbonate buffer | 16 h | MS(PEG)4 — short NHS-ester 100 mg - **$583.82** | Low; same moisture sensitivity as Round 1 |
| **10. Chamber Assembly** | Double-sided sticky tape, epoxy glue, streptavidin or neutravidin, T50 buffer | 60 min | Low cost for tape and epoxy glue. Streptavidin 10 mg - **$407.00** | Minimal |
| **TOTAL** | **-** | **~50 hours** | **$2455 AUD** | **Overall moderate risk** |

**From reference [5]**

**Table S2. Cost, timing and safety of rapid surface passivation for single molecule studies.**

| Step | Reagents | Time | Cost | Risk |
|---|---|---|---|---|
| **1. Piranha cleaning** | H2SO4 (conc.), H2O2 (30%), $NaHCO_3$ | 90 min | Sulfuric acid 500 mL - **$136.00** hydrogen peroxide 100 mL **- $125.00** | EXTREME — spontaneously heats >90C, violently oxidising, explosive with organics, fume hood required |
| **2. Rinse (post-piranha)** | Milli-Q water | 5 min | Negligible | Low |
| **4. NaOH etching** | NaOH solution (0.5 M) | 30 min | Sodium hydroxide solution 1L - **$44.10** | Moderate – corrosive base; gloves + eye protection |
| **5. Rinse (post-NaOH)** | Milli-Q water | 5 min | Negligible | Low |
| **7. Acetone soak** | HPLC-grade acetone | 20 min | Acetone 500 mL - **$72.50** | Moderate – flammable; no ignition sources |
| **8. Acetone rinse** | HPLC-grade acetone | 2 min | Same as above | Moderate – flammable |
| **9. Prepare PEG-silane solution** | mPEG-Sil-5000, biotin-PEG-Sil-5000, anhydrous DMSO | 20 min | mPEG-Sil-5000 1g - **$230.66** biotin-PEG-Sil-5000 1g - **$565.45** DMSO 50mL - **$143.00** | Moderate – DMSO is a penetrating solvent; anhydrous conditions critical; exclude moisture |
| **10. Pre-heat coverslip** | Glass chamber with desiccant (CaSO4) | 5 min | Negligible | Low – hot surface; use tongs |
| **11. PEG-silane incubation** | PEG-silane/DMSO solution (from step 11) | 15 min | Same as above | Moderate – hot DMSO; avoid evaporation; work quickly |
| **12. Rinse & dry** | Molecular biology grade water, dry N2 | 3 min | N/A | Low |
| **TOTAL** | **-** | **3.5 hours** | **$1316.71 AUD** | **Overall moderate risk** |

**From reference [6]**

**Table S3. Cost, timing and safety of the method developed in this work.**

| Step | Reagents | Time | Cost | Risk |
|---|---|---|---|---|
| **1. Lipid Mixing with chloroform** | Chloroform, DPhPC OR mPEG-DSPE | 30 min | Chloroform **$166** DPhPC **$900** mPEG-DSPE **$300** | Moderate – Chloroform is moderately toxic |
| **2. Lipid drying** | - | 16 h | - | None |
| **3. Lipid mixing into oil** | Silicone oil AR20 and Hexadecane | 45 min | Hexadecane **$300** Silicone Oil **$130** | Moderate – Hexadecane is toxic |
| **TOTAL** | **-** | **17 h** | **$1796 AUD** | Overal moderate risk |

Note: cost per-sample is estimated to be ~$3.30 AUD by extrapolating 200 samples possible with reagents listed.
